# Inhibition of IL34 unveils tissue-selectivity and is sufficient to reduce microglial proliferation in chronic neurodegeneration

**DOI:** 10.1101/2020.03.09.976118

**Authors:** Juliane Obst, Emilie Simon, Maria Martin-Estebane, Elena Pipi, Liana M. Barkwill, Ivette Gonzalez-Rivera, Fergus Buchanan, Alan R. Prescott, Dorte Faust, Simon Fox, Janet Brownlees, Debra Taylor, V. Hugh Perry, Hugh Nuthall, Peter J Atkinson, Eric Karran, C Routledge, Diego Gomez-Nicola

## Abstract

The proliferation and activation of microglia, the resident macrophages in the brain, is a hallmark of many neurodegenerative diseases such as Alzheimer’s disease (AD) and prion disease. Colony stimulating factor 1 receptor (CSF1R) is critically involved in regulating microglial proliferation, and CSF1R blocking strategies have been recently used to modulate microglia in neurodegenerative diseases. However, CSF1R is broadly expressed by many cellular types and the impact of its inhibition on the innate immune system is still unclear. CSF1R can be activated by two independent ligands, CSF1 and interleukin 34 (IL-34). Recently, it has been reported that microglia development and maintenance depend on IL-34 signalling. In this study, we evaluate the inhibition of IL-34 as a novel strategy to reduce microglial proliferation in neurodegenerative diseases, using the ME7 model of prion disease. Selective inhibition of IL-34 showed no effects on peripheral macrophages populations in healthy mice, avoiding the side effects observed after CSF1R inhibition on the systemic compartment. However, we observed a reduction in microglial proliferation after IL-34 inhibition in prion-diseased mice, indicating that microglia could be more specifically targeted by reducing IL-34 and that this ligand plays an important role in the modulation of microglia population during neurodegeneration. Overall, our results suggest that control of microglial response through IL-34 blockade could be a potential therapeutic approach in neurodegenerative diseases.

## Introduction

Neuroinflammation is a critical component of neurodegenerative diseases, including Alzheimer’s disease (AD), Parkinson’s disease (PD) or prion diseases. The neuroinflammatory process is characterised by a robust activation of the innate immune system, with an increase in the number of microglial cells associated with an activated and phagocytic phenotype ^1–3^. Experimental models of prion disease present several shared features of neurodegenerative diseases including protein misfolding, progressive synaptic degeneration followed by loss of neurons, microglial activation and production of inflammatory cytokines and chemokines^4^.

The contribution of local proliferation of microglia, regulated by the activation of the colony stimulating factor 1 receptor (CSF1R), has been shown as a disease-modifying mechanism during the progression of the ME7 prion model of progressive chronic neurodegeneration ^5^. Similarly, a prolonged inhibition of the tyrosine kinase activity of CSF1R blocks microglial proliferation and prevents synaptic degeneration, ameliorating disease progression, in the APP/PS1 model ^6^, the 3xTg model ^7^ and the 5xFAD model ^8,9^ of AD-like pathology. More recently, our group has validated this disease-modifying mechanism in the P301S mouse of tauopathy, demonstrating that inhibition of CSF1R is effective in repolarising microglia to a homeostatic phenotype, preventing neuronal degeneration ^10^. In recent years, the therapeutic potential of blocking antibodies and small molecule CSF1R kinase inhibitors has been demonstrated in inflammatory diseases, neurological disorders, bone diseases and cancer, with some candidates currently in clinical phase testing ^11–13^. However, the broader impact of this approach on the innate immune system remains unclear. CSF1R is expressed by cells of the myeloid lineage ^14^ and therefore it is anticipated that the inhibition of CSF1R would not only affect microglia, but also have potential on-target effects in circulating and tissue-resident myeloid populations, with a possible downstream immunosuppressive effect. CSF1R can be activated by two independent ligands with high affinity, the colony stimulating factor 1 (CSF-1) ^15^ and the recently identified interleukin 34 (IL-34) ^16^. A potential avenue to block this pathway more selectively is the specific modulation of the binding of its ligands, to increase tissue specificity and reduce side effects. Both ligands have been shown to promote microglial proliferation ^5^ but also show a differential temporal and spatial pattern of expression. Whereas CSF-1 is broadly expressed (spleen, lymph nodes, cortex, bone marrow, amongst others), the expression of IL-34 is restricted to a few tissues, predominantly produced in the skin and the brain, with minimal overlap with the CSF-1 expression pattern ^17,18^. Interestingly, mice lacking IL-34 (Il34^LacZ^) displayed a marked reduction of Langerhans cells in the skin and microglial cells in the brain, whereas other tissue macrophages were unaffected, showing that IL-34 specifically controls the development and maintenance of these populations ^19,20^.

Since IL-34 has been shown to be a tissue-restricted ligand of CSF1R, and it is crucial for the differentiation and maintenance of microglial cells in the brain, we aimed to investigate whether IL-34 inhibition could be used as a selective approach to reduce microglial proliferation during chronic neurodegeneration with minimal effects on other tissue-resident myeloid populations. Here, we first describe the effects of the selective inhibition of IL-34, compared to CSF1R inhibition, on different tissue-resident populations in healthy mice, supporting tissue-selectivity of IL-34. We also demonstrate that IL-34 inhibition decreases the proliferation of microglial cells in the ME7 prion model, showing that IL-34 is involved in the regulation of microglial proliferation and supporting that the inhibition of this cytokine could be used as a more selective therapeutic approach to modulate microglial proliferation in neurodegenerative diseases.

## Methods

### In vitro assessment of CSF1-R phosphorylation

The N13 murine microglia cell line ^21^ was cultured in Dulbecco’s modified Eagle’s medium (DMEM, Thermo Fisher Scientific), supplemented with 10% fetal bovine serum and 50U/mL penicillin/ streptomycin (Thermo Fisher Scientific). Cells were maintained in T75 flasks at 37 °C in a 5% CO_2_ humidified atmosphere. Cells were plated at a density of 2 × 10^5^ cells/cm^2^ in 6-well plates and cultured overnight to allow adherence. Cells were kept in serum-free medium for 4 hours prior to stimulation and then incubated with recombinant CSF-1 or IL-34 (R&D Systems) for the indicated time points, after which cells were immediately lysed in RIPA buffer (Thermo Fisher Scientific), supplemented with protease and phosphatase inhibitor cocktails (Roche, Thermo Fisher Scientific). Protein lysates were concentrated using Microcon-10kDa Centrifugal Filter Units (Merck Millipore), according to manufacturer’s instructions and protein concentration was determined using the Pierce BCA Protein Assay Kit (Thermo Fisher Scientific). Protein lysates were subjected to SDS-PAGE and Western blot.

### In vitro assessment of IL-34 neutralizing antibodies using CellTiter Glo

Mouse myelogenous leukemia (M-NFS-60) cells were M-CSF (R&D systems, 216-MC/CF) starved for 24 hours. In white clear bottom 96-well plates 10 μL IL-34 antibody (mouse monoclonal IgG2A (v1.1 manufactured by Genscript, Ma et al., 2012), rat monoclonal IgG2A (MAB5195, R&D Systems) and sheep polyclonal IgG (AF5195, R&D Systems) and 10 μL IL-34 stimulus (R&D systems, 5195-ML-CF) were incubated at 37°C for 30 minutes before 80 μL M-NFS-60 cells (10^3^ cells/well) were added. After two days incubation at 37°C cell viability was assessed using CellTiterGlo (Promega, G7570) following manufacturer’s instructions. 100 μL reconstituted CellTiterGlo was added per well, plates were shaken for 2 minutes and incubated at room temperature for 10 minutes before luminescence was read.

### Experimental model of prion disease and pharmacological treatments

C57BL/6J mice (Harlan laboratories), termed WT (wild type) throughout the manuscript, and *c-fms-*EGFP (macgreen) mice ^22^, characterized by eGFP expression under the *c-fms* (CSF1R) promoter, were bred and maintained in local facilities. Mice were housed in groups of 4 –10, under a 12 h light/12 h dark cycle at 21°C, with food and water *ad libitum*. To induce prion disease, 6 weeks old WT or macgreen mice were anesthetized with a ketamine/xylazine mixture (85 and 13 mg/kg), and 1 μL of ME7-derived (ME7 group) brain homogenate (10% w/v) was injected stereotaxically and bilaterally at the dorsal hippocampus, coordinates from bregma: anteroposterior, −2.0 mm; lateral, ±1.7 mm; depth, −1.6 mm. Naïve animals were used as controls. All procedures were performed in accordance with U.K. Home Office licensing. Group sizes were defined after performing power calculations, in order to achieve a significant difference of p<0.05, in light of a retrospective analysis of our previous published results, to reach a power between 0.80-0.90, depending on the specific experimental conditions. The effect size was calculated taking into consideration the strength of association between the variables, the sensitivity and the variation of any dependent variable. The calculations are the customary ones based on normal distributions.

For chronic treatment of healthy mice, rat monoclonal CSF1R-blocking antibodies (BE0213, Bio X Cell) and rat monoclonal IL-34 antibodies (MAB5195, R&D Systems) were diluted in PBS, pH 7.4 and administered by intraperitoneal injections 3 times a week for 3 weeks at a dose of 250 μg per injection. For chronic treatment in ME7 prion mice, mouse monoclonal IL-34 IgG2A ^23^ was administered biweekly for 4 weeks at a dose of 60mg/kg, starting 12 weeks after prion inoculation. For acute treatment in ME7 prion mice, 1 μg of mouse or human-specific IL-34 neutralizing antibody (sheep polyclonal IgG, AF5195 or AF5265, R&D Systems) was stereotaxically and bilaterally injected into the dorsal hippocampus, coordinates from bregma: anteroposterior, −2.0 mm; lateral, ±1.7 mm; depth, −1.6 mm, 12 weeks after induction of prion disease. Mice received daily intraperitoneal BrdU injections (7.5 mg/mL, 0.1 mL/10 g weight in sterile saline; Sigma-Aldrich) and were sacrificed one week after intracerebral antibody administration. Weight of the mice was monitored in all experiments, and no differences were observed between treated and untreated groups. All the experimental groups were randomised to avoid gender and cage effects, and all the experiments were performed double-blinded.

### Histology

Mice were terminally anesthetized with an overdose of sodium pentobarbital and transcardially perfused with 0.9% saline. Brain and peripheral organs (liver, kidney and spleen) were cut in serial sections (35μm thick) with a cryostat (Leica) and stored free-floating in cryoprotectant solution (30% sucrose, 30% ethylene glycol, 1% Polyvinyl-pyrrolidone (PVP-40) in 0.1M PB, pH 7.4) at −20°C. Immunohistochemistry of brain sections was performed as previously described ^5^. Briefly, sections were subjected to DNA denaturation with 2N HCl (30 min, 37°C), followed by incubation with 5% normal serum and 5% BSA in PBS to block nonspecific binding. After rinses with PBS-0.1% Tween 20 (PBST), sections were incubated overnight at 4°C with rabbit anti-Iba1 (Wako, 019-19741) and anti-BrdU (Biorad, MCA2060). After washes with PBST, sections were incubated with host-specific Alexa 488- and 568-conjugated secondary antibodies (Invitrogen). Brain sections and sections of peripheral organs from macgreen mice were counterstained with DAPI and mounted with Mowiol/DABCO (Sigma-Aldrich) mixture and imaged with a Leica DM5000B microscope coupled to a Leica DFC300FX camera.

### Analysis of skin-resident Langerhans cells

Ears were fixed in 4% paraformaldehyde overnight and then stored in PBS. The ears were split into dorsal and ventral halves and each pair was mounted on slide under coverslip mounted in Vectashield anti-fade mounting medium. For each half ear, 5 fields were chosen at random and 0.9-μm thick sections were collected on a LSM700 confocal microscope (Zeiss) using settings for eGFP fluorescence with a 40x objective. For each field a z-stack was taken to cover the full thickness of the Langerhans cells layer-typically 5-100 images depending on the flatness of the ear. Cell volume and number were measured using Volocity software (Quorum Technologies). Cells were identified as those with a GFP intensity >2SD from the mean image intensity. Non cellular objects <200 or >5000 μm^3^ were excluded. Objects with a long axis >100 μm were also excluded-this eliminated most auto-fluorescent hairs in the z stacks. Each image was checked manually to remove cell doublets and unexcluded hair profiles.

### Fluorescent activated cell sorting (FACS) analysis

Blood samples were harvested by cardiac puncture and collected in EDTA-coated tubes. Brain hemispheres were harvested in PBS 2%FCS 2mM EDTA (FACS buffer), mechanically triturated and enzymatically dissociated using the Neural Tissue Dissociation Kit (P) (Miltenyi). Samples were passed through a cell strainer of 70μm mesh (BD2 Falcon) with FACS buffer, and centrifuged at 500g for 10 min at 4°C. After the second wash, cells were re-suspended in 37% Percoll (GE Healthcare) and centrifuged at 500g for 30min at 18°C. The supernatant and myelin layers were discarded, and the cell pellet enriched with microglia/macrophages was resuspended in FACS buffer. Primary antibody labelling was performed for 1 hour at 4 °C, using the following antibodies: rat-anti-mouse CD11b (BD Biosciences, clone M1/70), rat-anti-mouse CD45 (Biolegend, clone 30-F11) and rat-anti-mouse Ly6C (BD Biosciences, clone AL-21) and Fixable Viability Dye eFluor™ 450 (eBioscience). Moreover, unstained cells and isotype-matched control samples were used to control for autofluorescence and/or non-specific binding of antibodies. Erythrocytes in blood samples were then lysed in RBC lysis buffer (eBioscience). Samples were run on a BD FACS Aria Flow cytometer. Data was analysed using FlowJo software.

### SDS-PAGE and Western blot

Frozen brain samples and peripheral organs were homogenized in T-PER™ Tissue Protein Extraction Reagent (Thermo Fisher), N13 cells were lysed in RIPA buffer (Thermo Scientific), supplemented with protease inhibitors (EASYpack, Roche) and phosphatase inhibitors (PhosSTOP, Roche). Homogenates were centrifuged at 13,000rpm and the supernatant was collected. Protein was quantified using BCA assay (Thermo Fisher) following the manufacturer’s instructions. 40 μg protein of N13 cell lysates was loaded to 7.5% Mini-PROTEAN® TGX Stain-Free™ Protein Gels (BioRad) and transferred to nitrocellulose membrane using the Trans-Blot® Turbo™ RTA Mini Transfer Kit (BioRad). After blocking with 5% BSA in TBS/0.1% Tween20, membranes were incubated with a combination of rabbit polyclonal antibodies against phospho-M-CSF receptor (Tyr546, Tyr708, Tyr723 and Tyr923, Cell signaling), phospho-AKT (Ser473, Cell signaling) or phospho-p44/42 MAPK (Erk1/2) (Thr202/Tyr204, Cell signaling) over night at 4°C. Membranes were washed in TBS and further incubated with an HRP-labelled anti-rabbit IgG (BioRad) for 2h at room temperature. Membranes were incubated with the SuperSignal™ West Pico Chemiluminescent Substrate (Thermo Fisher Scientific) and signal was detected on the ChemiDoc Imaging System (BioRad). Membranes were stripped using the Restore™ Western Blot Stripping Buffer (Thermo Fisher Scientific) and reprobed with mouse monoclonal CSF-1R antibody (D-8, Santa Cruz Biotechnology), anti-AKT (Cell signaling) or anti-ERK1/ERK2 antibody (9B3, abcam), followed by HRP-labelled anti-mouse or anti-rabbit IgG antibody (BioRad). Intensity of protein bands were quantified using Adobe Photoshop.

### ELISA

Nunc MaxiSorp™ flat-bottom 96-well plates (Thermo Scientific) were coated with F(ab’)2 fragment anti-rat or anti-mouse IgG (H+L) (Jackson ImmunoResearch) overnight. Plates were washed with PBS + 0.05% Tween20 and incubated with blocking buffer (PBS containing 0.05% Tween20 and 1% BSA) to block non-specific binding sites. Plasma samples or tissue lysates diluted in blocking buffer were incubated for 2h or overnight. A standard was generated using respective anti-IL-34 or anti-CSF1R antibodies that were used for in vivo treatment. After washing, plates were incubated with peroxidase-conjugated anti-rat or anti-mouse Fcγ subclass 2a-specific IgG (Jackson ImmunoResearch) for 2h, then washed and incubated with 1-Step™ Ultra TMB-ELISA Substrate Solution (Thermo Scientific). The reaction was stopped with 2N H_2_SO_4_ and the signal was measured on a plate reader at 450 nm.

CSF-1 and IL-34 in plasma or brain were measured by commercially available immunoassays (R&D systems), according to manufacturer’s instructions.

### Statistics

Data are shown as mean ± SEM and were analysed using the GraphPad Prism 6 software package (GraphPad Software), using two-way ANOVA with Tukey post-hoc test for multiple comparisons, Student’s t-test or one-way ANOVA followed by a Tukey post-hoc test for multiple comparisons, as indicated. Differences were considered significant at p < 0.05.

## Results

### IL-34 induces activation of the CSF1R signalling pathway and IL-34-mediated cell growth can be inhibited using neutralizing antibodies

To investigate whether IL-34 activates the CSF1R pathway in microglia, we stimulated a murine microglia cell line (N13) with either IL-34 or with CSF-1 and analysed tyrosine phosphorylation of the receptor and downstream signalling molecules ERK1/ERK2 and AKT. Stimulation with either IL-34 or CSF-1 leads to increased phosphorylation of CSF1R and downstream mediators, indicating that IL-34 binds to and activates the CSF1R pathway, triggering downstream signalling pathways related to survival and proliferation (Fig. 1A). IL-34-mediated growth of myelogenous leukemia cell line M-NFS-60 can be inhibited by three different IL-34 neutralizing antibodies, which were further used in this study and showed similar potencies (mouse monoclonal v1.1^24^: IC50 0.43 nM, rat monoclonal MAB5195: IC50 0.53 nM, sheep polyclonal AF5195: IC50 2.05 nM) (Fig. 1B), indicating that CSF1R-dependent signalling can be modulated by IL-34 inhibition.

**Figure 1:**
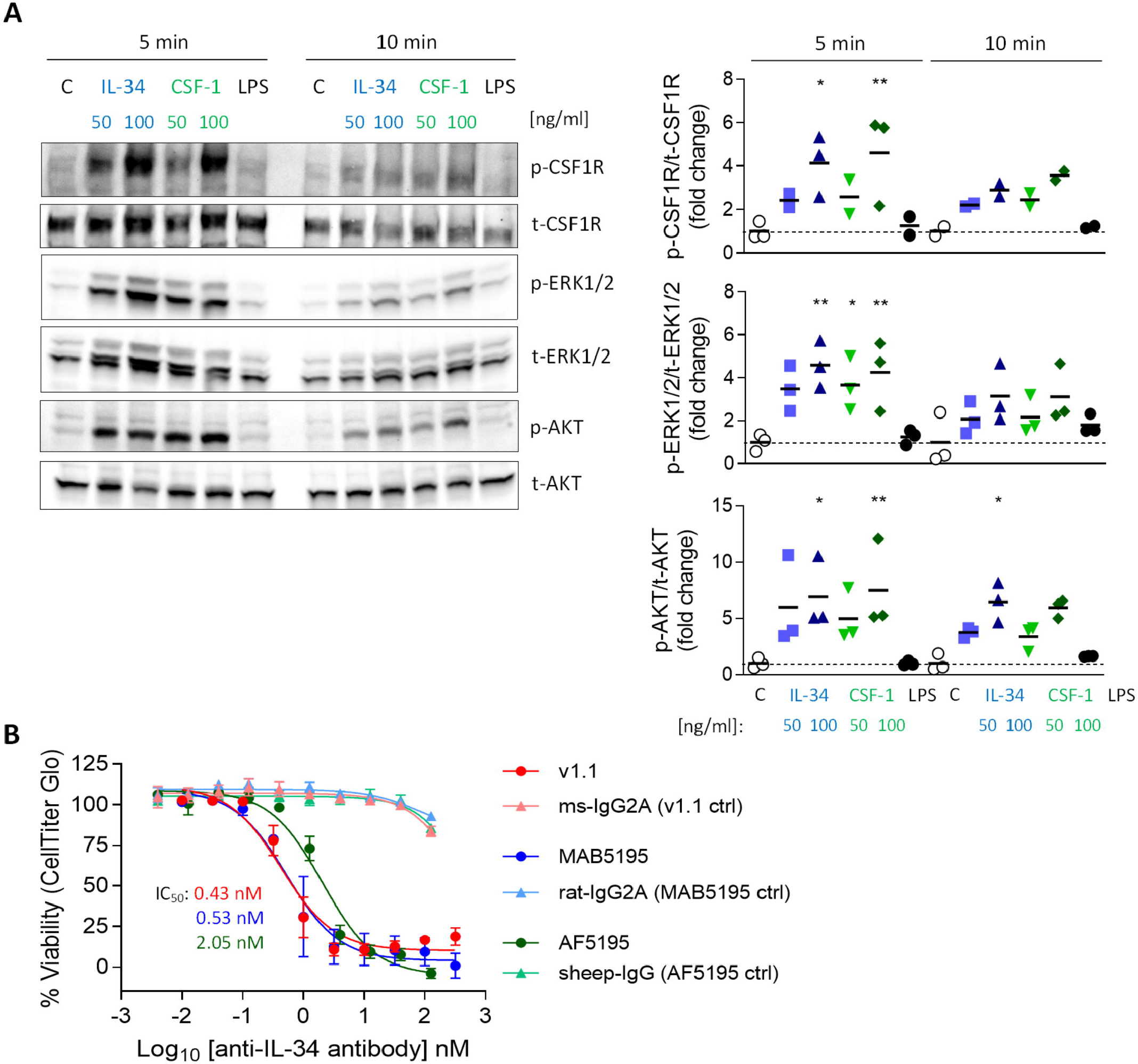
Activation of the CSF1R signalling pathway by IL-34 and CSF-1 and inhibition of IL-34-mediated cell growth using IL-34 neutralizing antibodies. [A] N13 microglia cell line was stimulated with IL-34 (50 or 100 ng/mL), CSF-1 (50 or 100 ng/mL) or LPS (1 μg/mL) for 5 or 10 minutes. Cell lysates were subjected to Western blotting which showed increased phosphorylation of CSF1R and downstream ERK1/2 and AKT after IL-34 and CSF-1 stimulation. [B] IL-34 neutralizing antibodies used in this study (mouse monoclonal IgG2A (v1.1, Ma et al., 2012), rat monoclonal IgG2A (MAB5195, R&D Systems) and sheep polyclonal IgG (AF5195, R&D Systems)) prevented IL-34-dependent cell growth of M-NFS-60 in a similar nanomolar range. n=3, data shown represent mean ± SEM, two-way ANOVA followed by Tukey’s multiple comparison test. * p < 0.05, ** p < 0.01, *** p < 0.001.

### CSF1R- but not IL-34 antibody-mediated inhibition leads to a reduction of CSF1R^+^/Ly6C^−^ blood monocytes

To determine the effect of CSF1R *vs.* IL-34 blockade on blood immune cells, macgreen mice were treated by intraperitoneal injections of CSF1R- or IL-34- neutralizing antibodies (monoclonal rat IgG2a, 250 μg per injection, 3 injections per week), vehicle (PBS) or rat IgG2a isotype for 3 weeks (Fig. 2A). The use of macgreen mice allows to monitor the abundance of CSF1R^+^ cells based on the *csf1r*-EGFP transgene reporter expression. Measuring antibody titer in the plasma following treatment using a rat IgG2a-specific ELISA showed comparable levels of antibody in anti-IL-34 and anti-CSF1R treated mice, while levels of isotype control were found to be significantly higher (Fig. 2B). Administration of CSF1R antibodies, but not IL-34 antibodies, increased CSF-1 levels in the plasma (Fig. 2C), which has been described as an indication of target engagement ^25^. Levels of IL-34 were found to be low in the plasma at baseline (approx. 30 times lower than CSF-1 levels) and were increased after CSF1R antibody treatment similar to CSF-1 levels, while IL-34 was undetectable after IL-34 antibody treatment (Fig. 2D). Flow cytometric analysis of blood immune cells demonstrated a significant decrease in CSF1R-positive monocytes after CSF1R antibody treatment, which was not significant after IL-34 antibody administration (Fig. 2E, F, for gating strategy see Suppl. Fig. 1). Further analysis of CSF1R^+^ subpopulations showed that the reduction in blood monocytes after CSF1R antibody treatment is largely due to an effect on cells with high CSF1R expression (eGFP^hi^) as opposed to cells with intermediate CSF1R expression (eGFP^int^, Fig. 2E). Furthermore, Ly6C^−^ monocytes were predominantly reduced after anti-CSF1R treatment, while Ly6C^+^ were not affected (Fig. 2F). Again, both Ly6C^+^ and Ly6C^−^ populations were not significantly reduced after IL-34 antibody administration. In contrast to blood immune cells, CSF1R^+^ cells in the bone marrow were not affected by either CSF1R or IL-34 antibody treatment, indicating that CSF1R is not required for the differentiation of myeloid progenitors in the marrow (Suppl. Fig. 2). Taken together, these results indicate that the number of CSF1R-positive monocytes in the blood depends on CSF1R, but is largely independent on IL-34.

**Figure 2:**
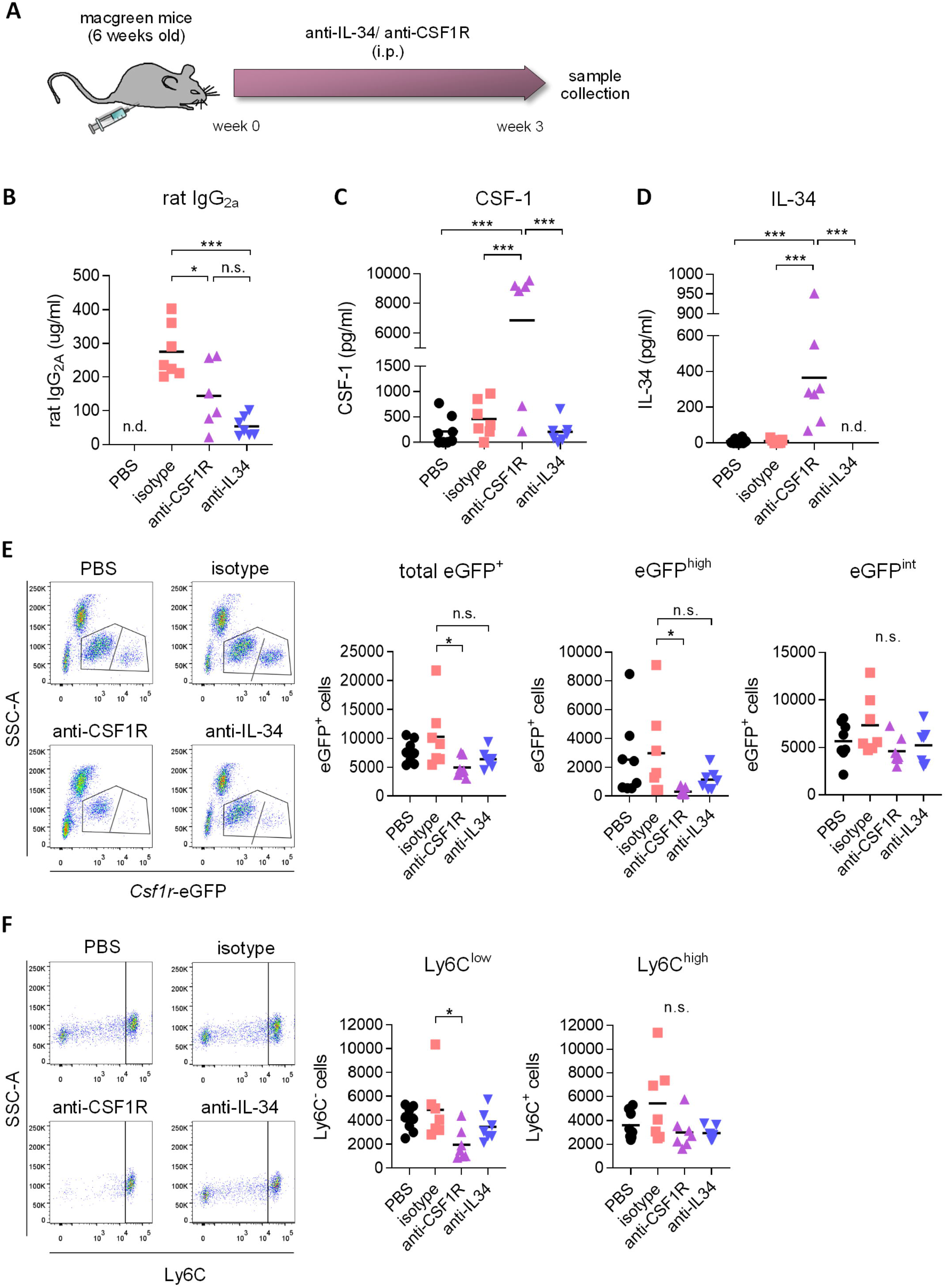
Effect of CSF1R- and IL-34 antibody treatment on blood immune cell compartment. [A] *c-fms*-eGFP (macgreen) mice were treated with anti-CSF1R or anti-IL-34 (both rat monoclonal IgG2A) by intraperitoneal injections of 250 μg antibody 3x a week for 3 weeks. [B-D] Levels of rat IgG2a, CSF-1 and IL-34 were measured in blood plasma after the treatment by ELISA, which showed increased IL-34 and CSF-1 levels after CSF1R-but not IL-34 blockade. [E] Flow cytometric analysis of CSF1R-positive cells in the blood of anti-CSF1R- and anti-IL-34- treated mice demonstrated a significant reduction after CSF1R-but not IL-34 antibody treatment. [F] Flow cytometry of Ly6C-positive and -negative subpopulations of CSF1R-expressing cells in the blood shows a reduction in Ly6C^low^ cells after anti-CSF1R treatment, while Ly6C^high^ cells were not affected. PBS n=8, isotype n=8, anti-CSF1R n=8, anti-IL-34 n=7, data shown represent mean ± SEM, two-way ANOVA followed by Tukey’s multiple comparison test. * p < 0.05, ** p < 0.01, *** p < 0.001.

### Systemic IL-34 blockade reduces epidermal Langerhans cells, but not macrophage populations in liver and kidney or microglia in the brain

We next aimed to determine the effect of IL-34 and CSF1R antibody treatment on different populations of tissue-resident macrophages. Measurement of rat IgG2a levels in liver, kidney, spleen and brain by ELISA showed equal distribution of IL-34- and CSF1R-neutralizing antibodies in each organ, with a distribution between different organs from highest to lowest as follows: spleen > kidney > liver > brain (Suppl. Fig. 3). Administration of IL-34 neutralizing antibodies for 3 weeks did not change the number of CSF1R-positive macrophages in the liver and in the kidney (Fig. 3A). In contrast treatment with a CSF1R blocking antibody lead to a pronounced reduction of macrophages in both organs, demonstrating a 41% reduction in liver-resident macrophages and a 85% reduction of macrophages in the kidney (Fig. 3A). Skin-resident CSF1R^+^Langerhans cells, were significantly decreased after treatment with either IL-34- or CSF1R blocking antibodies (Fig. 3A). This indicates that skin-resident Langerhans cells depend on IL-34- mediated signalling through CSF1R, while macrophages in liver and kidney depend on IL-34 independent CSF1R signalling.

**Figure 3:**
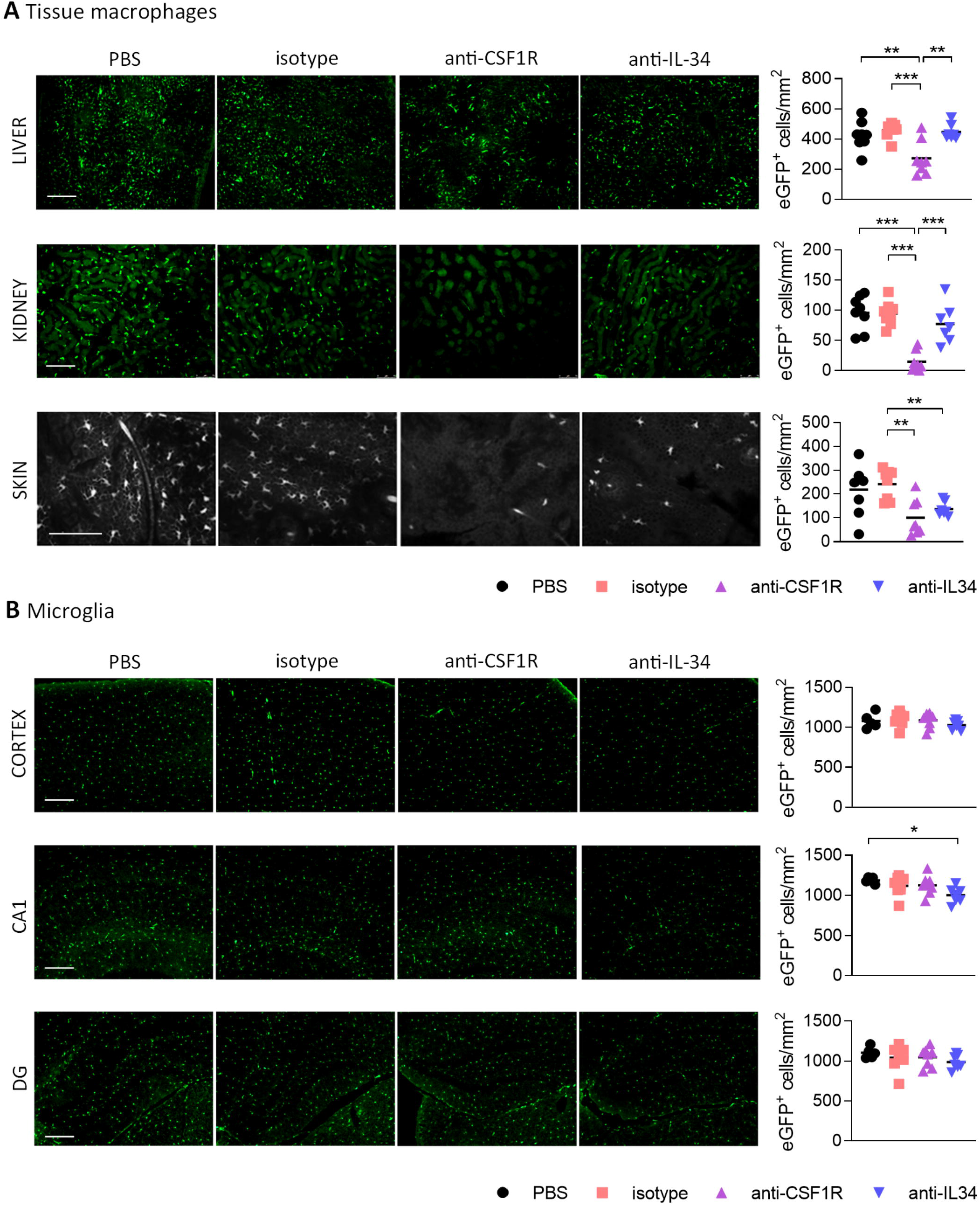
Effect of CSF1R- and IL-34 antibody treatment on tissue macrophages. [A] Histological analysis of CSF1R-positive cells in liver and kidney shows a reduction after anti-CSF1R treatment, but not after anti-IL-34 treatment. Skin Langerhans cells were significantly reduced after both CSF1R – and IL-34 blockade. [B] Microglia in cortex, hippocampal CA1 and dentate gyrus were not majorly affected by CSF1R or IL-34 antibody administration. Scale bar 100 μm. PBS n=8, isotype n=8, anti-CSF1R n=8, anti-IL-34 n=7, data shown represent mean ± SEM, two-way ANOVA followed by Tukey’s multiple comparison test. * p < 0.05, ** p < 0.01, *** p < 0.001.

To find out whether IL-34 antibody treatment affects microglia number in the brain we quantified the number of CSF1R^+^ cells in the cerebral cortex, hippocampal CA1 and dentate gyrus. Peripheral administration of CSF1R or IL-34 blocking antibodies for 3 weeks did not overtly affect the number of microglia in the brain, with only a small reduction in the CA1 region of the hippocampus observed after anti-IL-34 administration. (Fig. 3B). Since previous reports showed that CSF1R inhibition using small molecule inhibitors leads to a reduction in microglia ^5^, it is likely that we did not reach optimal antibody penetration into the brain with the administered dose of antibody to achieve sufficient target engagement.

### Chronic systemic administration of IL-34 blocking antibody lacks sufficient central target engagement, not modifying the microglial population in ME7 prion mice

We next aimed to investigate whether chronic systemic IL-34 antibody treatment would affect microglia numbers in the ME7 prion mice, a model of neurodegeneration which is characterized by a pronounced expansion of the microglia population ^5^. Based on our previous observation showing a lack of effect in the brain with 250 μg antibody per injection (~10 mg/kg, Fig. 3B), we increased the administered dose to 60 mg/kg per injection (Fig. 4A). After 4 weeks of biweekly intraperitoneal injections of the antibody in prion diseased mice, microglia populations were analysed by flow cytometry and histology. While ME7 prion mice showed increased numbers of microglia compared to naïve animals, there was no difference in microglia numbers in brains of anti-IL-34 treated animals compared to control-treatment (Fig. 4B, C). Levels of isotype and anti-IL-34 antibody were detectable in plasma and brain using a mouse-IgG2a specific ELISA, which revealed higher levels of isotype compared to IL-34 antibody in both compartments, with a brain/plasma ratio of 0.165 for the isotype and 0.141 for the IL-34 antibody (Fig. 4D). CSF-1 levels in the brain were significantly increased in prion mice compared to naïve mice, but unaffected by the anti-IL34 antibody treatment (Fig. 4E). IL-34 levels were around 300 times higher in the naïve brain than CSF-1 levels, but not changed in the context of prion disease or after IL-34 antibody treatment (Fig. 4E). In order to determine whether the absence of an effect of IL-34 antibody treatment on microglia numbers could be due to insufficient target engagement, we developed an ELISA to capture mouse IgG2a from brain lysates, followed by detection of IL-34 molecules bound to captured IgG2a. Using this assay we detected bound IL-34 exclusively in brain lysates of mice treated with IL-34 antibody, but not with isotype or PBS (Fig. 4F). The percentage of IL-34 bound to IL-34 antibody showed a dose dependent increase after 2 injections in healthy mice, demonstrating increased target engagement with increasing doses of IL-34 antibody (Fig. 4F). However ME7 prion mice treated for 4 weeks with biweekly injections of 60 mg/kg showed that only approximately 13% of total IL-34 was bound to the antibody, suggesting a low degree of target engagement (Fig. 4F).

**Figure 4:**
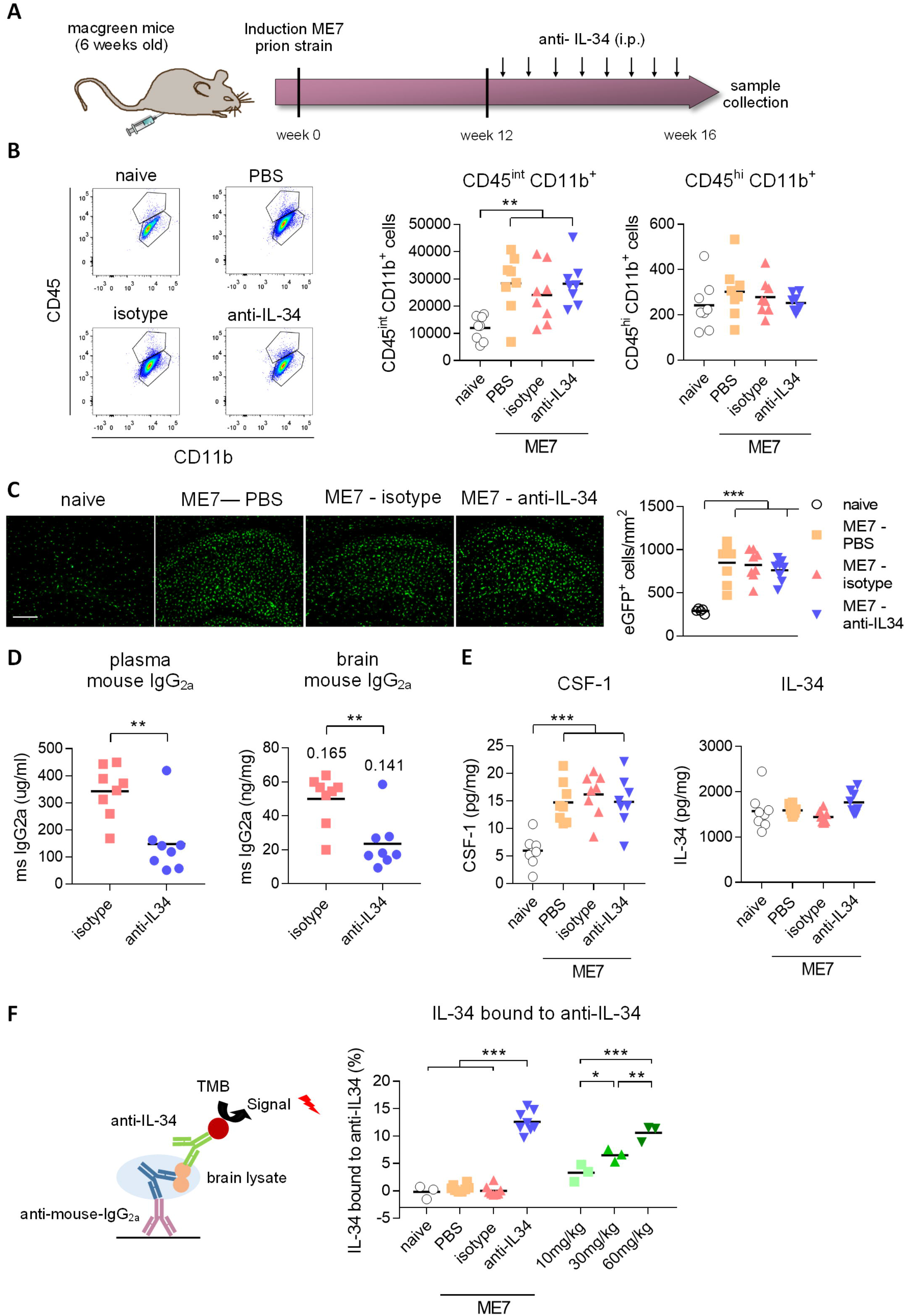
Peripheral IL-34 antibody injections in ME7 prion mice did not affect microglia numbers. [A] Macgreen mice infected with prion disease were treated with anti-IL-34 (mouse monoclonal IgG2A) by intraperitoneal injections at a dose of 60 mg/kg twice a week for 4 weeks. [B] Flow cytometric analysis of CSF1R-positive cells in the brain of anti-IL-34- treated mice showed no effect on number of CD45^int^ or CD45^hi^ microglia. [C] Histological analysis of CSF1R-positive cells in the cortical brain shows unchanged microglial numbers after anti-IL-34 treatment. [D] Levels of mouse IgG2a were measured in blood plasma and brain lysates by ELISA, which showed higher levels of isotype than anti-IL-34. Values for tissue/plasma ratio are indicated above bars. [E] CSF-1 and IL-34 were measured in brain lysates, showing no alterations of IL-34 and CSF-1 levels after anti-IL-34 administration. [F] IL-34 bound to anti-IL34 in brain lysates were detected by ELISA, by coating plates with anti-mouse IgG2a to capture IL-34 antibodies present in brain lysates and detecting IL-34 bound to IL-34 antibodies with an IL-34 specific detection antibody (R&D systems). Levels of IL-34 bound to IL-34 antibodies were less than 15% from total IL-34 levels. Scale bar 100 μm. n = 8 per group, data shown represent mean ± SEM, two-way ANOVA followed by Tukey’s multiple comparison test. * p < 0.05, ** p < 0.01, *** p < 0.001.

### Intracerebral administration of IL-34 blocking antibodies reduces microglia proliferation in ME7 prion mice

Since we did not achieve a significant degree of target engagement in the brain with peripheral antibody administration, we aimed to determine whether administering the antibody directly into the brain of prion mice could affect microglia numbers. IL-34 antibodies were stereotactically injected into the hippocampus 12 weeks after induction of prion disease, a timepoint of pronounced microglia proliferation ^5^, and brains were collected one week after injection (Fig. 5A). Detection of the proliferation marker BrdU and microglia marker Iba1 showed increased microglial proliferation and microglia numbers in the hippocampus of prion mice compared to naïve mice, while injection of a mouse-specific IL-34 neutralizing antibody significantly reduced microglia proliferation by about 50% (Fig. 5B). Administration of a human-specific IL-34 antibody did not have an effect on microglia proliferation, possibly due to reduced homology with mouse IL-34 (71%^26^). The reduction of microglial proliferation however did not result in a reduction of total microglia numbers after IL-34 antibody administration at the analysed timepoint, probably due to the acute and transient nature of the intervention (Fig. 5B). The finding that microglia proliferation was locally reduced after direct intracerebral injection provides proof-of-concept that IL-34 is involved in regulating microglia proliferation in the context of chronic neurodegeneration and that IL-34 blockade could be used therapeutically to reduce microglia proliferation.

**Figure 5:**
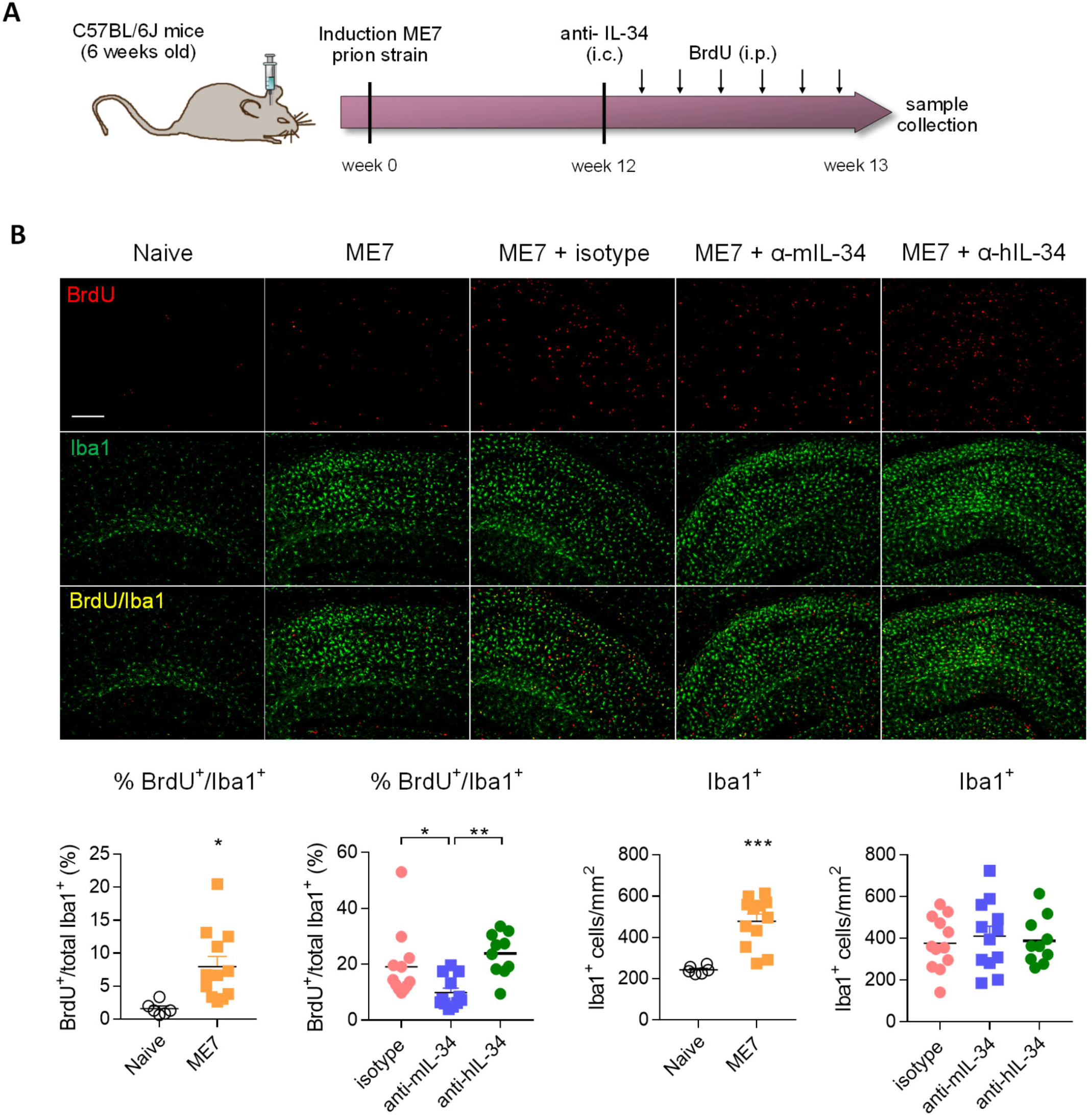
Intracerebral IL-34 antibody administration resulted in reduced microglia proliferation in ME7 prion mice. [A] Mice infected with prion disease received a single intracerebral injection of mouse- or human-specific anti-IL-34 (sheep polyclonal IgG) and brains were analysed one week later. [B] Histological analysis of BrdU/Iba1-positive microglial cells in the cortex showed a reduction after treatment with a mouse-, but not with a human-specific antibody. Scale bar 100 μm. Naïve n=6, ME7 n=12, ME7 + isotype n=10, ME7 + anti-mIL-34 n=8, ME7 + anti-hIL-34 n=7, data shown represent mean ± SEM, two-way ANOVA followed by Tukey’s multiple comparison test. * p < 0.05, ** p < 0.01, *** p < 0.001.

## Discussion

In this study we explored the effect of inhibition of IL-34 on blood monocytes, systemic tissue macrophages and microglia in health and neurodegenerative disease, modelled by a murine model of prion disease. IL-34 is a tissue-specific ligand of the CSF-1 receptor predominantly expressed in the brain and in the skin and has been shown to be crucial for the development and survival of microglia and epidermal Langerhans cells^19,20^. We showed here that inhibition of IL-34 does not affect monocyte and macrophage populations in many peripheral tissues, with an exception of skin-resident Langerhans cells. Even though we did not achieve sufficient target engagement in the brain after peripheral administration of IL-34 blocking antibodies, we observed a local reduction of microglia proliferation when injecting IL-34 antibodies directly into the brain of mice infected with prion disease, indicating that IL-34 is involved in driving proliferation of microglia in the context of neurodegenerative disease.

IL-34 and CSF-1 were shown to activate the CSF1R signalling cascade in a similar manner^27^ and have an overall similar effect on human monocyte differentiation *in vitro* as shown by transcriptional profiling and pathway analysis^28^. We have also found that the activation pattern of CSF1R and downstream pathway components AKT and ERK1/2 induced by IL-34 resembles the one induced by CSF-1 in microglia. Although IL-34 and CSF-1 seem to similarly affect CSF1R activation and macrophage differentiation, they have distinct tissue expression pattern with limited spatial overlap^27^. In accordance with previous reports which showed that IL-34 is more widely expressed in the brain than CSF-1^17,19,20^, we observed that overall levels of IL-34 protein are approximately 300 times higher than CSF-1, highlighting its predominant role in the brain. However, we also observed that IL-34 levels did not further increase in the brain of prion-diseased mice.

It was demonstrated previously that CSF1R inhibition using tyrosine kinase inhibitors can be used as a strategy to decrease microglia proliferation in neurodegenerative disease models, which led to beneficial effects such as reduced neuronal loss and behavioural deficits in mouse models of prion disease^5^, tau pathology^10^ and Aβ pathology^8^,^7^. Long-term CSF1R inhibition could potentially increase risk of infections and lead to disturbance of tissue homeostasis due to the reduction of CSF1R-dependent macrophage populations in multiple organs. As such, CSF1R antagonism leads to higher susceptibility of mice to lethal West Nile virus infection and lack of virologic control in both brain and periphery^29^. However, a CSF1R tyrosine kinase inhibitor showed a good safety and tolerability profile in patients with rheumatoid arthritis over a course of 3 months, causing only minor side effects related to compromised Kupffer cell function^30^, however the long-term consequences of CSF1R pharmacological inhibition are not well understood. A potential way to circumvent affecting multiple macrophage populations is to target IL-34, which is predominantly expressed in the brain. We aimed to evaluate whether inhibition of this more tissue-restricted ligand of CSF1R would cause a reduction in microglia proliferation without having a major impact on peripheral myeloid cell populations. In support of this strategy, the administration of recombinant IL-34 into the brain caused locally increased microglia proliferation similar to CSF-1^5^, showing that IL-34 can induce proliferation of microglia in the brain. In IL-34^lacZ/lacZ^ mice which lack IL-34, microglial numbers are strongly reduced in many brain regions such as cortex, hippocampus and striatum^19,20^, indicating that IL-34 is at least partially responsible for maintenance of the population. Targeting IL-34 using an antibody-based approach, we showed that specific inhibition of IL-34 was sufficient to reduce microglia proliferation present in prion pathology, at least when IL-34 was inhibited over a short period of time by administration of IL-34 antibodies directly into the brain. This finding suggests that IL-34 is partially involved in regulating microglia proliferation in neurodegenerative disease. However, it proved to be challenging in our study to target brain-intrinsic IL-34 using systemically administered neutralizing antibodies probably due to their poor brain penetrance which prevented sufficient antibody titres to efficiently neutralize biological function of IL-34 in the brain. Although two recent reports showed a reduction in microglial numbers after peripheral administration of IL-34-specific monoclonal antibodies at high doses^31,32^, we did not observe a modulation of microglial numbers after chronic systemic antibody treatment in healthy mice or mice infected with prion disease. We found IL-34 to be approximately 300 times higher in the brain than CSF-1, while in the blood it was nearly absent. This high abundance of IL-34 in the brain emphasizes the difficulty in efficiently targeting this cytokine with neutralizing antibodies Similarly, peripheral administration of CSF1R blocking antibodies did not lead to a reduction in microglia in cortex, dentate gyrus and CA1. The fact that blocking CSF1R using small molecule inhibitors resulted in pronounced depletion of microglia in several mouse models^5,7,8,10^, further indicates that the strategy applied in this study using blocking antibodies is not favourable. Lin *et al.* and Easley-Neal *et al.*, provide first proof that manipulation of IL-34 can be used to modify the microglia population in the gray matter of most brain regions^32^ and that this approach might be relevant in the context of inflammatory diseases and cancer^31^. We extend these findings by showing that in a model of neurodegenerative disease, which is characterized by a pronounced expansion of the microglia population predominantly in the hippocampus, inhibition of IL-34 leads to reduced microglial proliferation. In order to provide further proof-of-concept that IL-34 inhibition can be used to tackle microglial proliferation in neurodegenerative disease over a longer amount of time, it is inevitable to apply other strategies which offer an improved brain penetrance profile.

Chronic inhibition of IL-34 did not affect the abundance of blood monocytes or tissue-resident macrophage populations in liver and kidney which were sensitive to CSF1R blockade, suggesting IL-34- independent mechanisms of survival, most likely through CSF-1. In line with this, a natural mutation in the *Csf-1* gene *(op/op)* caused a reduction in macrophages in most tissues of the body^12^, and long-term treatment with a CSF1R-blocking antibody lead to a complete depletion of Kupffer cells in the liver and prevented the development of nonclassical Ly6C^+^ monocytes in the blood ^33^, which we likewise observed. It was previously shown, that genetic deficiency of IL-34 resulted in decreased numbers of microglia in most brain regions, while there was no effect on myeloid cells in blood and bone marrow, Kupffer cells in the liver, lung alveolar macrophages and dendritic cells in the lung and spleen^19^. Similarly, a specific impact of IL-34 blockade on Langerhans cell homeostasis, but not on liver, intestine and kidney macrophages after IL-34 antibody administration has been recently shown^31^. We have also observed an effect on Langerhans cells, which were reduced after both CSF1R- and IL-34 antibody treatment, confirming a role of IL-34 in regulating their survival as well as the efficacy of anti-IL-34 antibodies. Overall, sensitivity of macrophage populations to IL-34 inhibition seems to be defined by the spatial expression pattern of IL-34, which rarely overlaps with CSF-1 expression. Thus, myeloid cells located in regions dominated by IL-34 expression can be targeted by inhibition of IL-34, which is restricted to fewer organs, potentially reducing unwanted side effects caused by therapeutic intervention targeting the CSF1R pathway. We propose that IL-34 inhibition could be a viable strategy to decrease proliferation of microglia in the context of neurodegenerative disease, with restricted impact on peripheral myeloid cells.

## Acknowledgements

We thank Georgina Dawes for technical assistance. We thank the Southampton Flow Cytometry Facility for technical advice, and the Biomedical Research Facility for assistance with animal breeding and maintenance. The research was funded by the Alzheimer’s Research UK Dementia Consortium and Medical Research Council (MR/P024572/1).

## Supplementary figure legends

**Suppl. figure 1:** Gating strategy used for flow cytometric analysis of brain, blood and bone marrow after CSF1R- and IL-34 antibody treatment.

**Suppl. figure 2**: Effect of CSF1R- and IL-34 antibody treatment on bone marrow. [A] Flow cytometric analysis of CSF1R-positive cells in the bone marrow of anti-CSF1R- and anti-IL-34- treated mice did not result in significant alterations. [B] Flow cytometry of Ly6C-positive and -negative subpopulations of CSF1R-expressing cells in the bone marrow did not reveal any significant changes due to the treatment. PBS n=8, isotype n=8, anti-CSF1R n=8, anti-IL-34 n=7, data shown represent mean ± SEM, two-way ANOVA followed by Tukey’s multiple comparison test. * p < 0.05, ** p < 0.01, *** p < 0.001.

**Suppl. figure 3:** Distribution of CSF1R- and IL-34 antibodies in peripheral organs and brain. Levels of rat IgG2a were measured in tissue lysates of brain, liver, kidney and spleen after the treatment by ELISA, showing no significant differences between anti-CSF1R and anti-IL-34 in individual organs. Brain: PBS n=8, isotype n=8, anti-CSF1R n=8, anti-IL-34 n=7, liver/kidney/spleen: PBS n=4, isotype n=4, anti-CSF1R n=3, anti-IL-34 n=4, data shown represent mean ± SEM, two-way ANOVA followed by Tukey’s multiple comparison test. * p < 0.05, ** p < 0.01, *** p < 0.001.

