## Supplementary figures and images for "Inhibition of IL34 unveils tissue-selectivity and is sufficient to reduce microglial proliferation in chronic neurodegeneration"

### Supplemental Figure 1

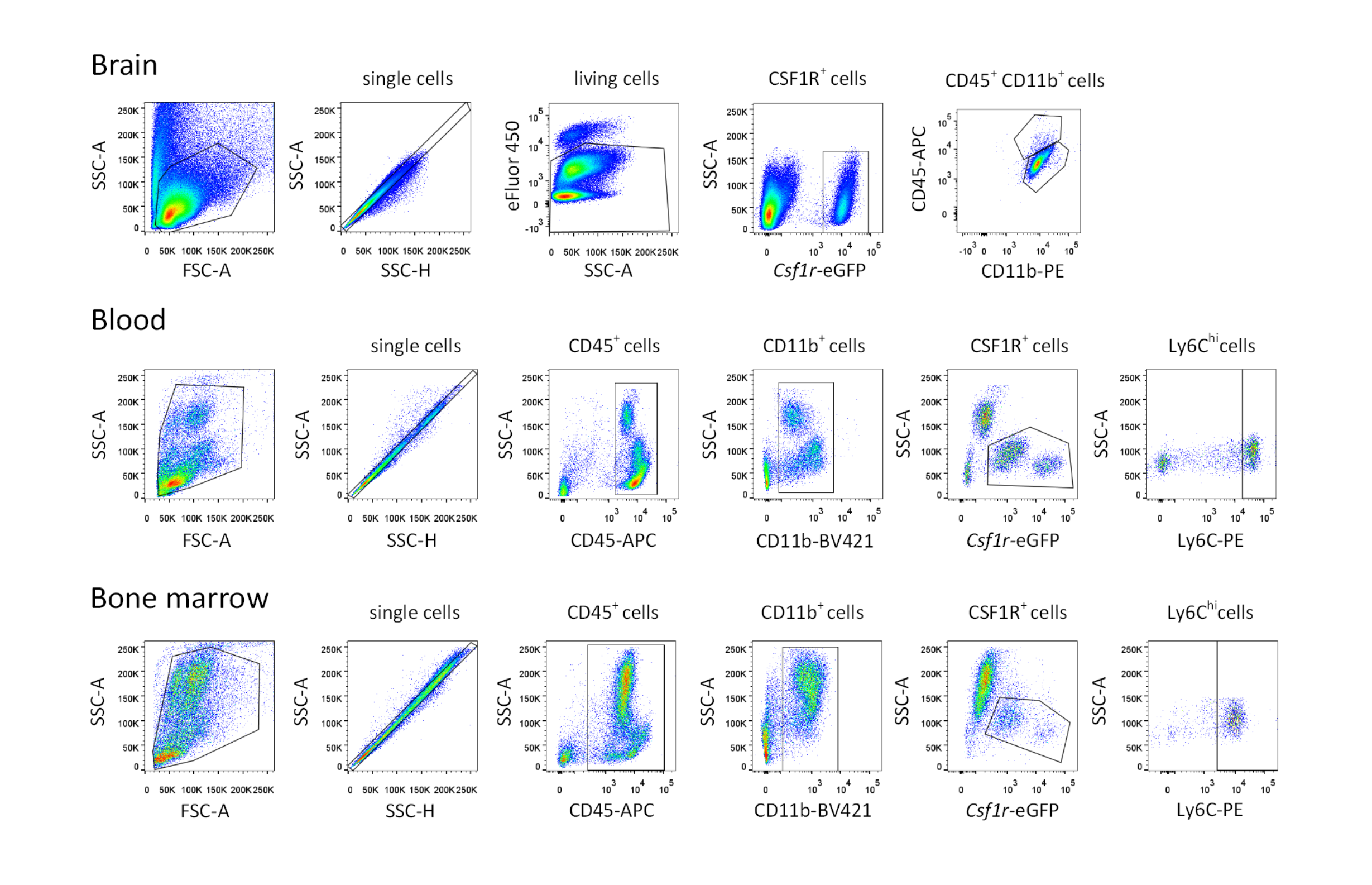

### Supplemental Figure 2

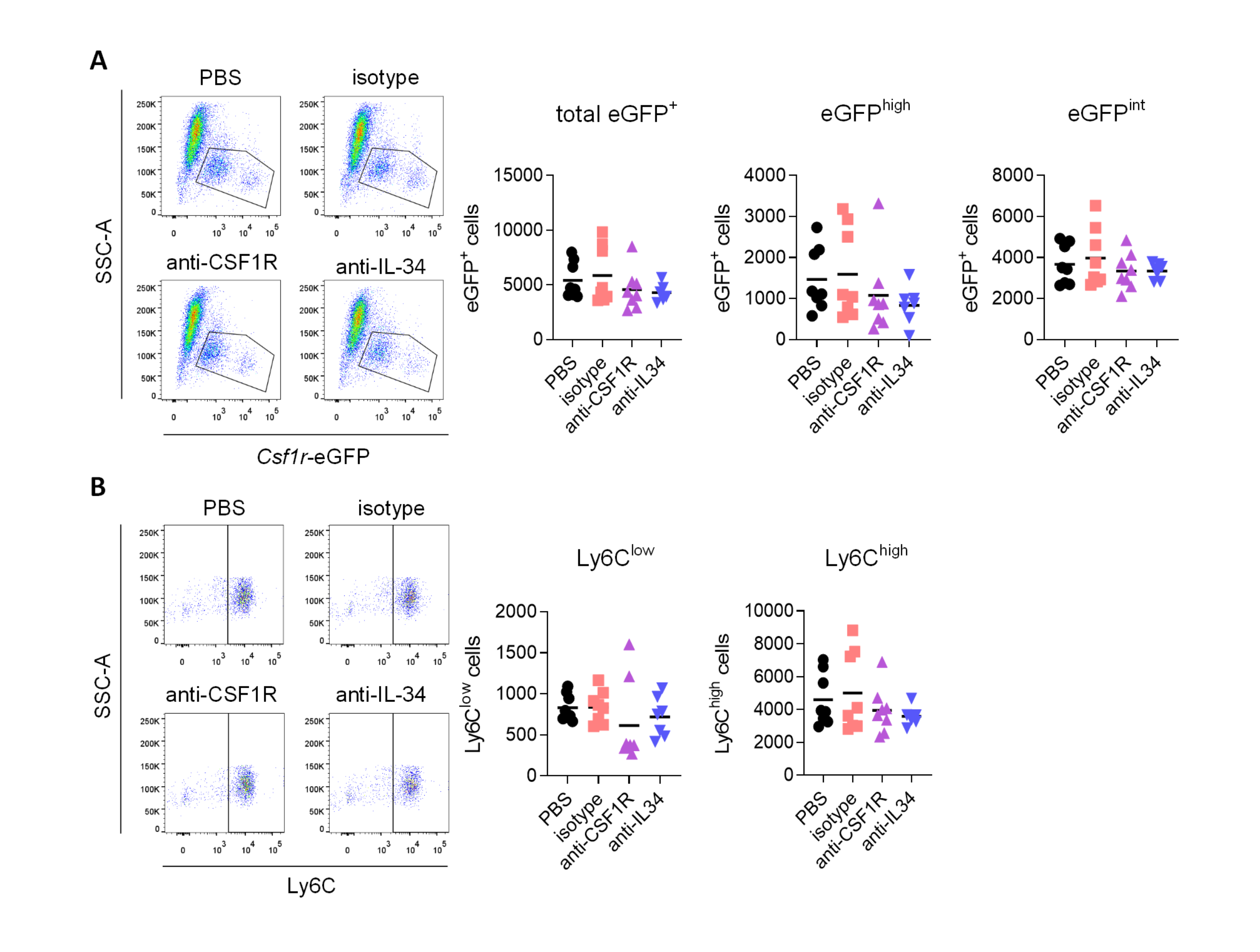

### Supplemental Figure 3

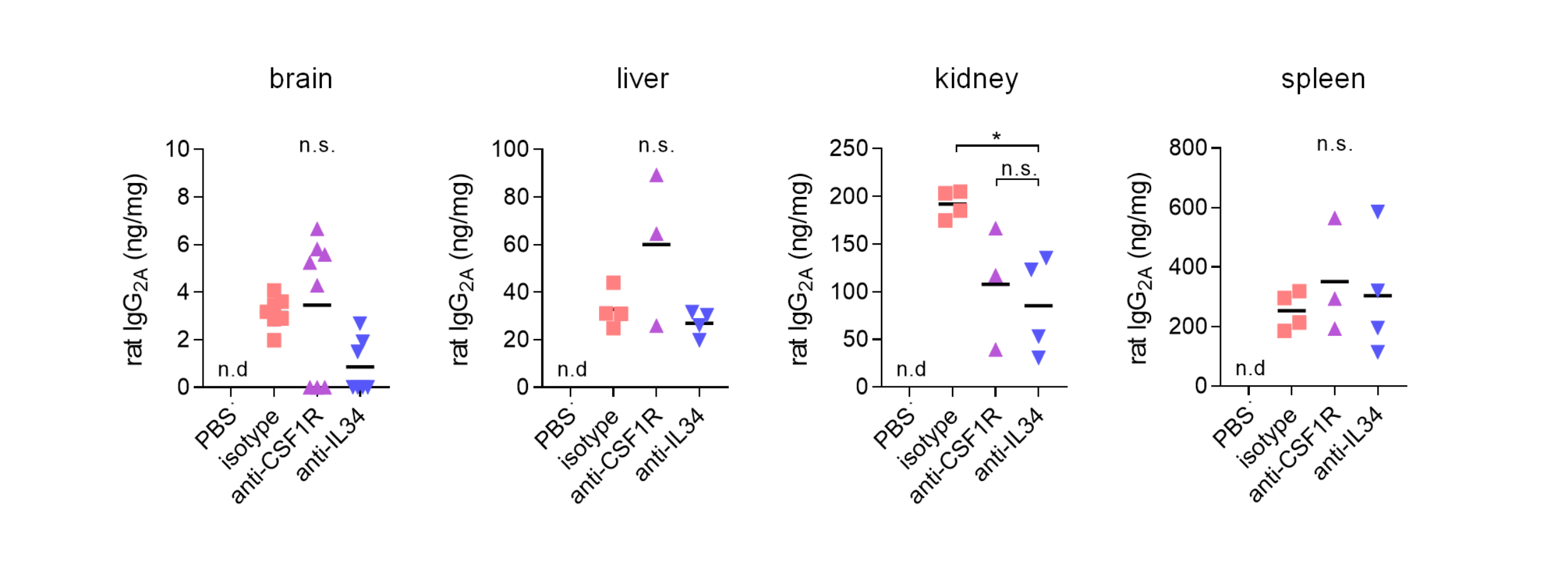
